# The Small Non-coding RNA miR-16-1-3p Hampers Cancer Stem Cell Self-renewal and Invasiveness, Boosting Chemosensitivity by Adjusting TGF-β1 Signaling via MDM2/p53 Axis in Human Osteosarcoma

**DOI:** 10.1101/2025.11.19.689220

**Authors:** Wenyu Xue, Yuzhe Wang, A.V. Smirnova, P.A. Malakhov, Margarita Pustovalova, Denis V. Kuzmin, Sergey Leonov

## Abstract

The TGF-β signaling pathway has both tumor-suppressing and metastasis-promoting effects in cancer. However, the molecular determinants governing this switch remain unclear. Here, we explored the miR-16-1-3p/MDM2/p53 axis as a critical conductor of the TGF-β-Smad pathway in osteosarcoma.

Although miR-16-1-3p overexpression by itself markedly reduces proliferative and clonogenic potential of U2OS cells, when paired with TGF-β treatment, it significantly increases arrest cells in G1 phase and nearly extinguishing the growth capability of these cells. MiR-16-1-3p overexpression inhibited TGF-induced actin remodeling and EMT featuring, significantly decreasing vimentin levels. TGF-β enhances both 2D and 3D migration, but miR-16-1-3p overexpression, alone or with TGF-β, strongly counteracts its pro-migratory effects. MiR-16-1-3p restored p53 stability by targeting MDM2, redirecting TGF-β-Smad signaling toward p21 activation and proliferation inhibition while attenuating its EMT-promoting capacity. Administration of TGF-β together with miR-16-1-3p dramatically increases the sensitivity of wild-type U2OS cells to cisplatin, exceeding that of TGF-β therapy alone by more than an order of magnitude. Administering TGF-β and miR-16-1-3p together significantly reduces the tumor nodule volume and Ki67 expression, while effectively eradicates metastases in the chicken chorioallantoic membrane (CAM) in vivo model.

For the first time, our research demonstrates that miR-16-1-3p shifts TGF-β1 signaling from a facilitator of metastasis to a promoter of anti-growth effects through MDM2 inhibition and p53 stabilization, effectively reducing the self-renewal and invasiveness of cancer stem cells in human osteosarcoma model. This process preserves TGF-β’s tumor-suppressive role while limiting its associated cancer risks.

## Introduction

Transforming growth factor-β (TGF-β) plays a dual role in cancer development, serving as a tumor suppressor in the early phases but shifting to a tumor promoter in later stages of the disease. The tumor-suppressing effects, mainly through Smad-induced cell cycle arrest and apoptosis, contrast sharply with its pro-cancer roles, such as promoting epithelial-mesenchymal transition (EMT), invasion, and metastasis.^1,2^ This functional dichotomy poses a major therapeutic challenge: inhibiting TGF-β’s tumor-promoting arm risks eliminating its intrinsic tumor-suppressive capacity.^3^

Osteosarcoma, the most common primary malignant bone tumor in adolescents and young adults, reflects this dilemma.^4,5^ While some studies suggest that TGF-β slows down osteosarcoma cell growth and increases response to treatment, ^6,7^ others indicate it aids in the spread of cancer.^8–10^

These opposing effects have been observed across independent experimental systems. No study has yet systematically explored both the tumor-suppressive and tumor-promoting effects of TGF-β signaling in a unified cellular context, nor has it identified the molecular factors that determine its functional outcomes. Critically, it remains unclear what governs the shift between pro-tumorigenic and anti-tumorigenic TGF-β signaling in osteosarcoma. The intracellular conditions that bias this pathway toward either outcome are poorly defined. Although Smad-interacting cofactors and post-translational modifiers are linked to other cancers, their specific roles in osteosarcoma are not yet known.^11^

Emerging evidence indicates that p53 is considered a key determinant of the functional direction of TGF-β signaling.^12^ MDM2, an E3 ubiquitin ligase best known as a negative regulator of p53, is overexpressed in aggressive and metastatic osteosarcoma.^13,14^ Studies have found that MDM2 enhances migration induced by TGF-β, and p53 is known to regulate TGF-β signaling in some contexts.^12,15,16^ However, it remains unclear whether the MDM2–p53 axis influences the balance between TGF-β’s tumor-suppressive and tumor-promoting outputs in osteosarcoma.^15^ MDM2 not only blocks p53 but may also boost TGF-β-induced migration and an epithelial-mesenchymal transition (EMT), increasing its cancer-promoting effects.^17,18^ Discovering this regulatory mechanism would clarify TGF-β’s dual role and allow for strategies to keep its benefits.

Based on this, we turned our attention to upstream regulators of TGF-β1/MDM2 signaling. The role of non-coding RNAs (ncRNAs) in modulating TGF-β signaling is complex and context-dependent. Analyzing how ncRNAs affect the TGF-β1 pathway is important for designing effective therapies that manage tumor growth and metastasis in cancer. MicroRNAs (miRNAs) are ncRNAs that control gene expression by binding to complementary sequences in the 3′ untranslated regions (3′UTRs) of target genes. miRNAs play crucial roles in tumor development and are gaining attention as new therapeutic targets.^19^ miR-16-1-3p, a passenger strand of the miR-16-1, has been shown to exert tumor-suppressive functions ^20^ and increasing sensitivity to DNA-damaging stress ^21^across multiple cancers. However, whether miR-16-1-3p, can exert a similar effect in TGF-β signaling context in osteosarcoma is unknown.

Our hypothesis posits that the role of miR-16-1-3p in TGF-β signaling entails diminishing MDM2 levels while activating p53, instigating a transformative change in signaling outcomes: moving from promoting migration and EMT phenotype to successfully inhibiting tumors and inducing cell cycle arrest. In this research, we thoroughly validated the regulatory axis within an integrated osteosarcoma in vitro and in vivo model, with the purpose of shedding light on the inherent mechanisms driving the dual roles of TGF-β and delivering new perspectives for improving TGF-β-targeted therapeutic approaches.

## Results

### MDM2 attenuates the anti-proliferative effect of TGF-**β**1 and enhances its pro-migratory impact

U2OS cells, derived from human osteosarcoma, are an early-stage epithelial-like cell line retaining wild-type p53, and serve as an ideal model to investigate the dual functions of TGF-β1 and its regulatory mechanisms. We explored the effect of MDM2 on TGF-β1’s biological effects in U2OS cells with active p53. Cells were treated with TGF-β1 (5 ng/mL), TGF-β1 plus MDM2 overexpression, and a control, and we analyzed proliferation and migration.

In colony formation assays, TGF-β1 significantly inhibited colony number, exhibiting a canonical anti-proliferative effect (Figure□1A). MDM2 overexpression not only reversed the suppression but boosted colony growth, showing it disrupts TGF-β1’s tumor-suppressive function. The SRB proliferation assays confirmed TGF-β1’s effect in reducing cell growth, while MDM2 co-treatment significantly enhanced proliferation (Figure□1B).

Both 2D collective (wound healing assay) and 3D confined (Transwell assay) migration demonstrated that TGF-β1 significantly increased cell motility. MDM2 overexpression significantly enhanced wound closure and invasion through the membrane (Figures□1C–D).

Overall, these findings indicate that MDM2, when p53 is functional, undermines TGF-β1’s anti-growth impact and increases its migratory effects, steering TGF-β1 signaling toward promoting tumors. This suggests MDM2 as a key molecular switch governing the functional output of TGF-β1 signaling.

### miR-16-1-3p boosts the anti-proliferative capabilities of TGF-**β**1 and mitigates its pro-migratory effects via MDM2 targeting

A dual-luciferase reporter assay was performed in Hela cells to determine if miR-16-1-3p targets MDM2 and influences TGF-β1 functions. Co-transfection of miR-16-1-3p mimics with a luciferase reporter carrying the MDM2 3’ UTR greatly decreased luciferase activity. No changes were seen with the empty control vector, showing that miR-16-1-3p can directly target MDM2 (Fig. 2A). Western blot analysis in U2OS cells further confirmed that MDM2 protein levels were markedly decreased following miR-16-1-3p overexpression (Fig. 2B).

**Figure 1.**
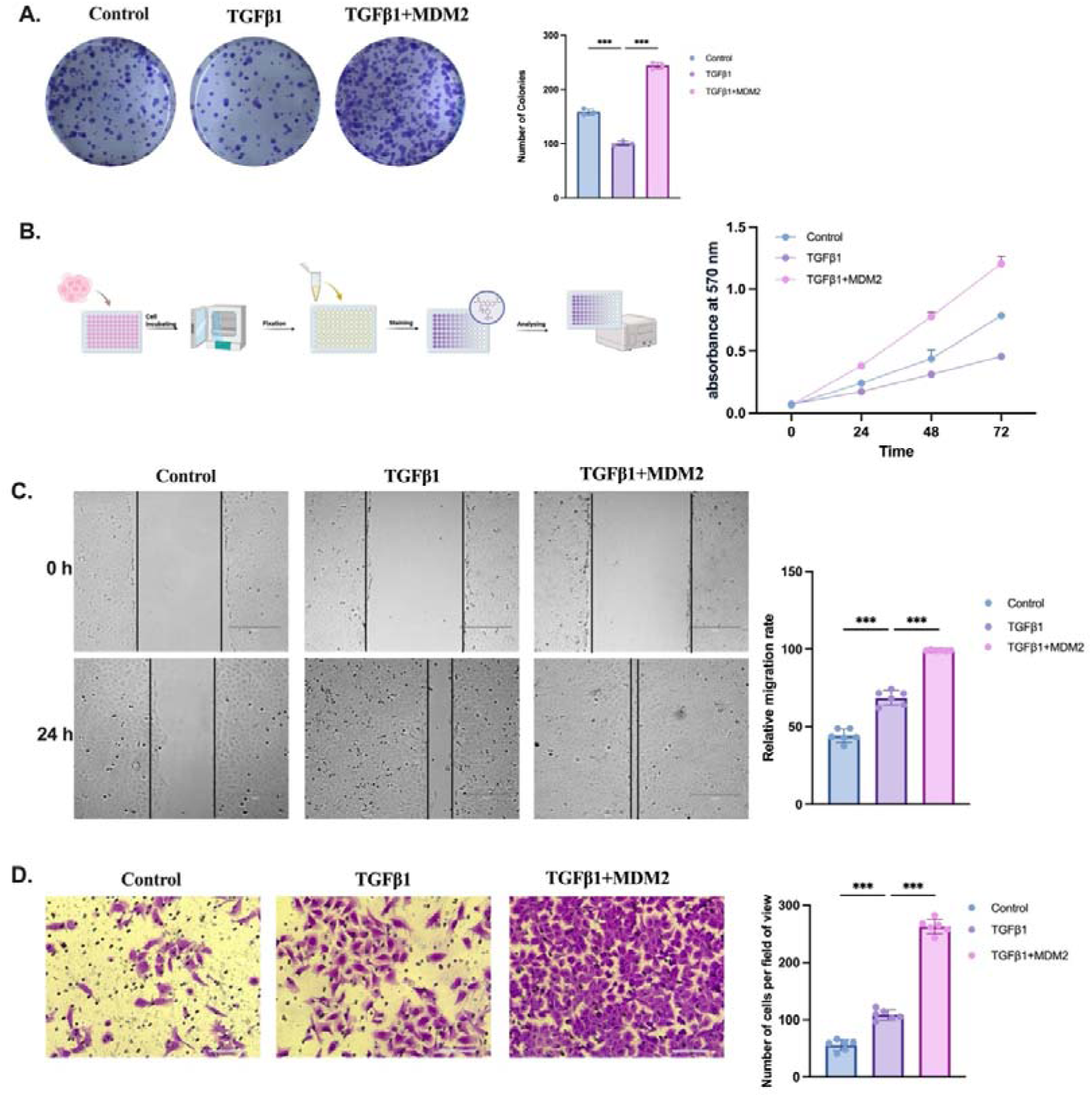
MDM2 augments the TGF-β1-induced effects on U2OS cells. (A) Colony formation assay of U2OS cells treated with 5ng/ml TGF-β1, with or without MDM2 overexpression. Images of crystal violet-stained cell colonies (left) and the number of colonies quantified (right) at 10 days after seeding. (B) SRB assay of cell proliferation: total cell mass (absorbance at 570 nm) was measured at 0, 24, 48, and 72 h after seeding 5000 cells/well in 96-well plates.(C) Wound healing assay: images of wound area (left) taken at 0h and 24h after “scratch” made, and its quantification (right). Scale bar= 300 um. (D) Transwell assay: Images at 20x magnification display crystal violet-stained cells on the outer membrane of the inner chamber (left). It also depicts the cell migration rate 24 hours after seeding 100000 cells/well in the upper Boyden chamber of a 24-well Transwell insert (right). Scale bar= 50 um.

**Figure 2.**
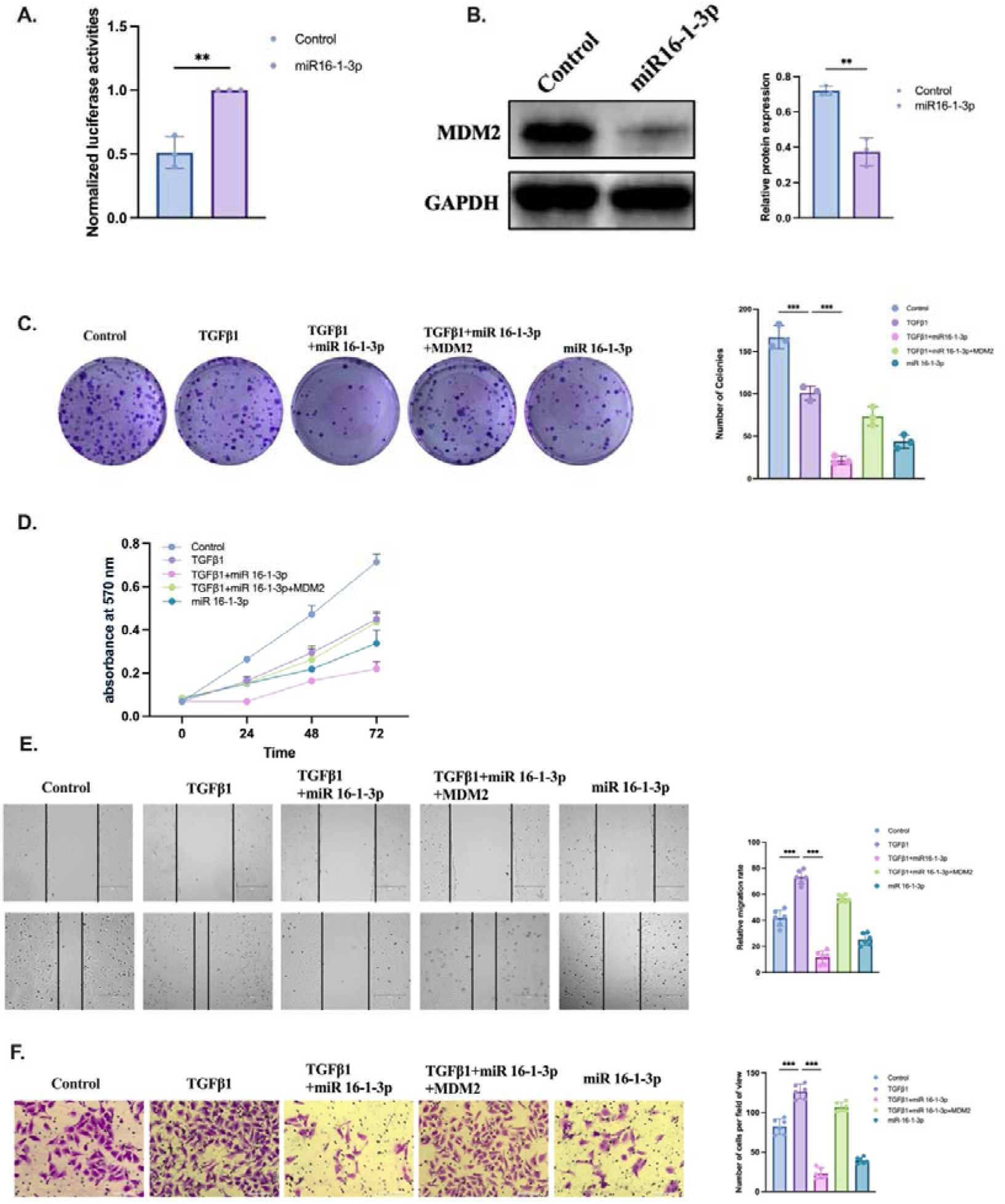
Influences of miR-16-1-3p mimics on the TGF-β1-mediated effects in U2OS cells. (A) Dual-luciferase reporter assay in Hela cells co-transfected with miR-16-1-3p mimics (50□nM) and a luciferase construct containing the wild-type MDM2 3′UTR, confirming direct targeting. (B) Western blot showing MDM2 protein levels in U2OS cells 24□h after transfection with miR-16-1-3p mimics (50□nM) or Scramble control. (C) Colony formation assay in U2OS cells co-transfected with miR-16-1-3p mimics (50□nM) or Scramble control and MDM2 overexpression plasmid or empty vector. After 24□h, 500 cells/well were seeded in 6-well plates and treated with TGF-β1 (5□ng/mL) during colony formation. (D) SRB assay of cell proliferation: total cell mass (absorbance at 570 nm) was measured at 0, 24, 48, and 72 h after seeding 5000 cells/well in 96-well plates.(E) Wound healing assay: images of wound area (left) taken at 0h and 24h after “scratch” made, and its quantification (right). Scale bar= 300 um. (F) Transwell assay: Images at 20x magnification display crystal violet-stained cells on the outer membrane of the inner chamber (left). The cell migration rate 24 hours after seeding 1000 cells/cm² in the upper Boyden chamber (right). Scale bar= 50 um.

U2OS cells treated with TGF-β1 were transfected with miR-16-1-3p mimics, both alone and in combination with MDM2, to determine the significance of this regulatory interaction. Addition of miR-16-1-3p mimics significantly potentiates TGF-β1-induced attenuation of colony formation, while MDM2 overexpression restores the clonogenic capacity of U2OS cells (Fig. 2C). SRB assays showed that adding miR-16-1-3p mimics boosts the TGF-β1-induced reduction in cell proliferation, which MDM2 co-expression can partially reverse (Fig. 2D).

Higher levels of miR-16-1-3p mimics mitigate TGF-β1’s effectiveness in promoting cell migration in wound healing and Transwell tests, leading to slower wound closure and fewer migrating cells. These suppressive effects on migration were partially reversed upon MDM2 restoration (Fig. 2E–F).

These results reveal that the increase of miR-16-1-3p decreases MDM2 protein expression, enhancing the anti-proliferative effects of TGF-β1 while opposing its migration-inducing abilities. Restoration of MDM2 expression abrogates these effects, underscoring the functional importance of the miR-16-1-3p–MDM2 axis in modulating TGF-β1 signaling output.

### miR-16-1-3p reconfigures TGF-**β**1 downstream signaling by targeting MDM2, regulating cell cycle progression and EMT phenotype

TGF-β1 initiates its signaling cascade by first attaching to the TGF-β1 type II receptor (TβRII), which subsequently recruits the TGF-β1 type I receptor (TβRI or Alk5) to establish a heterodimeric serine/threonine kinase complex. This active heterodimeric complex initiates the transphosphorylation of TβRI, allowing it to directly phosphorylate two carboxyterminal serines on Smad2 and Smad3, which ultimately activates these transcription factors^22^. Accordingly, TGF-β1 stimulation significantly activated the Smad signaling pathway, indicated by higher phosphorylation of Smad2/3 in U2OS cells (Figure 3A).

**Figure 3.**
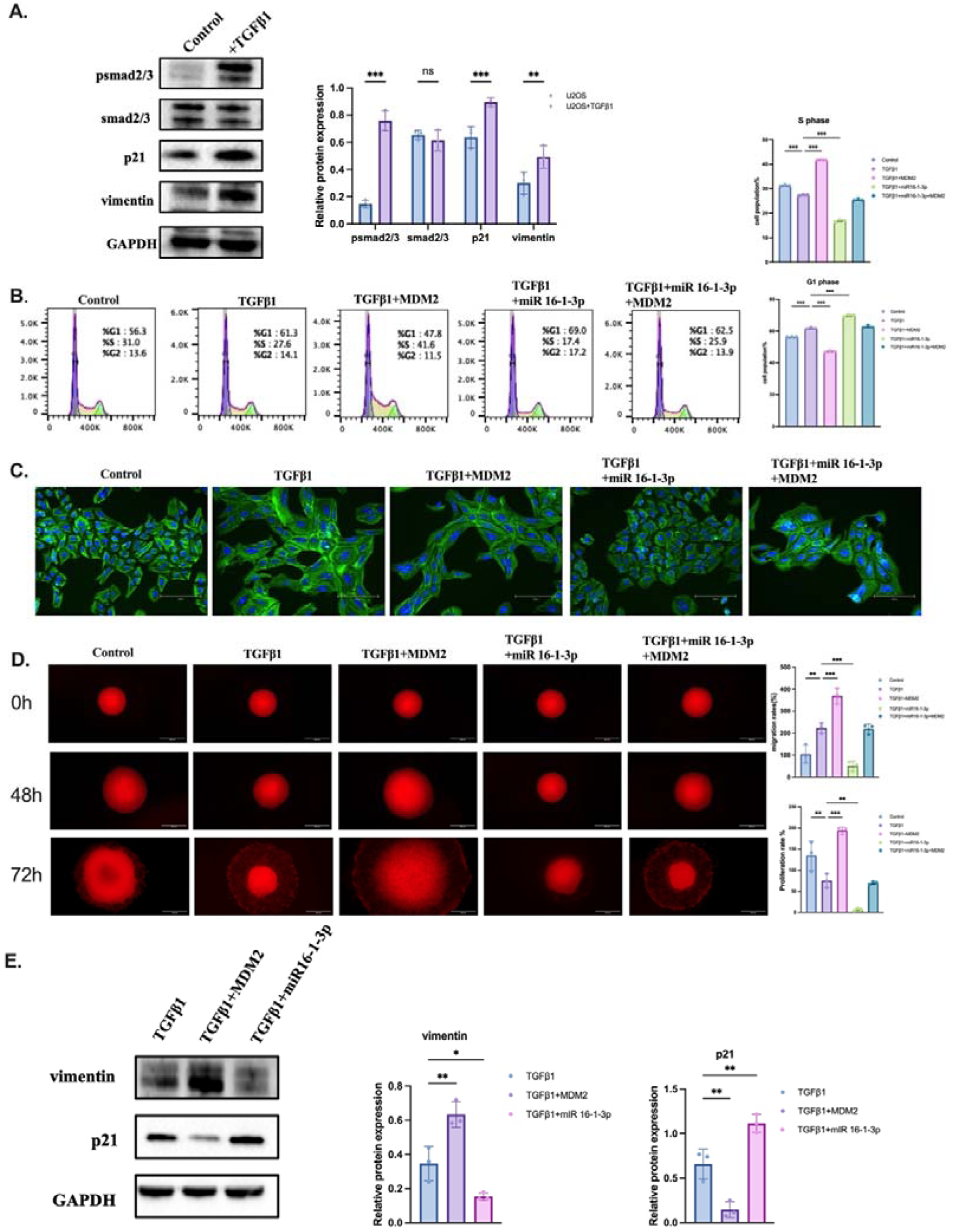
Impact of miR-16-1-3p on TGF-β1 downstream signaling. (A) Western blot analysis of pSmad2/3, total Smad2/3, p21 and vimentin expression in U2OS cells treated with 5ng/ml TGF-β1 for 24h (left). Quantification of protein expression (right). GAPDH expression was used for a protein loading control. U2OS exposed to 5ng/ml TGF-β1 and 50 nM of miR-16-1-3p mimics after 24 h with or without MDM2 overexpression: (B) Flow cytometry analysis of cell cycle distribution. Quantification of TGF-β1 promoted S-phase (upper right), and G1 arrest in U2OS cells (bottom right). (C) TGF-β1 promoted cytoskeletal reorganization and EMT-like phenotype indicated by phalloidin fluorescence staining of F-actin remodeling. Scale bar= 150 um. (D) 3D tumorsphere assay embedded in collagen matrix. Spheroids were first formed in agarose for 48h, then embedded in collagen for another 24h to evaluate invasion. Invasion was evaluated after 24h of serum deprivation. Scale bar= 200 um. Quantification of migration rates (upper right) and proliferation rates (bottom right). (E) Western blot of vimentin and p21 expressions (left). Quantification of protein expression (right). GAPDH expression was used for a protein loading control.

The pairing of Smad2 or Smad3 with Smad4 leads to their translocation into the nucleus, where these activated complexes engage with Smad-binding elements (SBEs) located in the promoters of many genes ^23,24^. Certain targeted genes, including p21, regulate processes that slow growth and may trigger cell death under certain conditions ^25^. We found that TGF-β1 activation raised levels of p21, a key cell cycle inhibitor, and vimentin, a mesenchymal cell marker, in U2OS cells (Figure 3A). This indicates that TGF-β1 can induce both cell cycle arrest and epithelial–mesenchymal transition at the same time.

To delineate the functional implications of these molecular changes, we first analyzed cell cycle progression by flow cytometry. TGF-β1 treatment markedly increased the proportion of cells in the G1 phase, indicating an anti-proliferative effect (Figure 3B). Overexpression of MDM2 promoted the G1–S transition and reduced G1 phase accumulation. In contrast, transfection with miR-16-1-3p mimic further enhanced the G1 arrest induced by TGF-β1. Notably, this effect was partially reversed by co-transfection with an MDM2 overexpression plasmid, confirming that miR-16-1-3p modulates TGF-β1-induced cell cycle arrest through targeting MDM2 (Figure 3B).Our analysis of the cytoskeleton showed that activation of TGFβ/Smad signaling caused U2OS cells to adopt a mesenchymal-like shape, featuring elongated bodies and strong actin stress fibers highlighted by phalloidin staining (Figure 3C). These features were further accentuated by MDM2 overexpression. Conversely, miR-16-1-3p treatment markedly reversed the morphological transition, restoring epithelial-like cell arrangement, consistent with its EMT-suppressive role (Figure 3C).

Cancer stem cells (CSCs) have been identified as a primary factor contributing to the recurrence of cancer. Growing tumorspheres is essential for evaluating cancer stem cells’ (CSCs) self-renewal and for isolating CSCs from a larger group of cancer cells □. The 3D tumorsphere model can also be used to study how TGF-β1 influences cell invasiveness. By embedding spheroids in a collagen 3D matrix, researchers can study the impact of TGF-β1 signaling on cancer cell invasion in a setup that resembles a natural tumor.

CSC self-renewal and invasive behavior were evaluated using a tumorsphere culture model. During the early culture phase, TGF-β1 treatment resulted in a reduction in tumorsphere diameter, whereas under subsequent collagen matrix conditions, TGF-β1 promoted outward invasive growth of the spheres. In contrast, miR-16-1-3p inhibited both tumorsphere growth and invasion, while MDM2 overexpression enhanced these phenotypes (Figure 3D).

To further confirm the molecular basis of these phenotypic changes, we performed Western blot analysis across different treatment groups. miR-16-1-3p increased p21 and decreased vimentin expression under TGF-β1 stimulation, whereas MDM2 overexpression suppressed p21 and upregulated vimentin, mirroring the observed trends in cell cycle progression and EMT (Figure 3E).

These findings show that miR-16-1-3p inhibits both CSCs self-renewal and invasiveness via adjustment of TGF-β1 signaling by targeting MDM2, boosting its anti-proliferative effects through p21 and limiting its pro-migratory effects by suppressing vimentin. Hence, MDM2 is key in TGF-β1 signaling, influencing whether cells enter growth arrest or progress to metastasis.

### miR-16-1-3p redirects the functional TGF-**β**1 signaling output via the MDM2–p53 axis

Results from Western blot analysis highlighted a crucial relationship between TGF-β1 signaling direction and the MDM2–p53 axis status. Overexpression of miR-16-1-3p effectively downregulated MDM2 and restored p53 protein levels. Enhanced p53 expression altered pSmad signaling, with higher p21 levels and lower vimentin levels. This suggests that TGF-β1 signaling combined with p53 promotes anti-proliferative effects and reduces EMT traits (Figure 4A). TGF-β1 stimulation induced p21 expression at both the transcriptional (Figure 4B) and protein levels(Figure 4A). Nevertheless, qPCR confirmed that enhanced miR-16-1-3p significantly raises p21 expression in both mRNA (Figure 4B) and protein (Figure 4A) compared to TGF-β1, but MDM2 overexpression prevents this enhancement. This indicate that MDM2 reduces p21 levels, possibly through complex post-translational mechanisms, whereas miR-16-1-3p boosts p21 in a more direct way.

**Figure 4.**
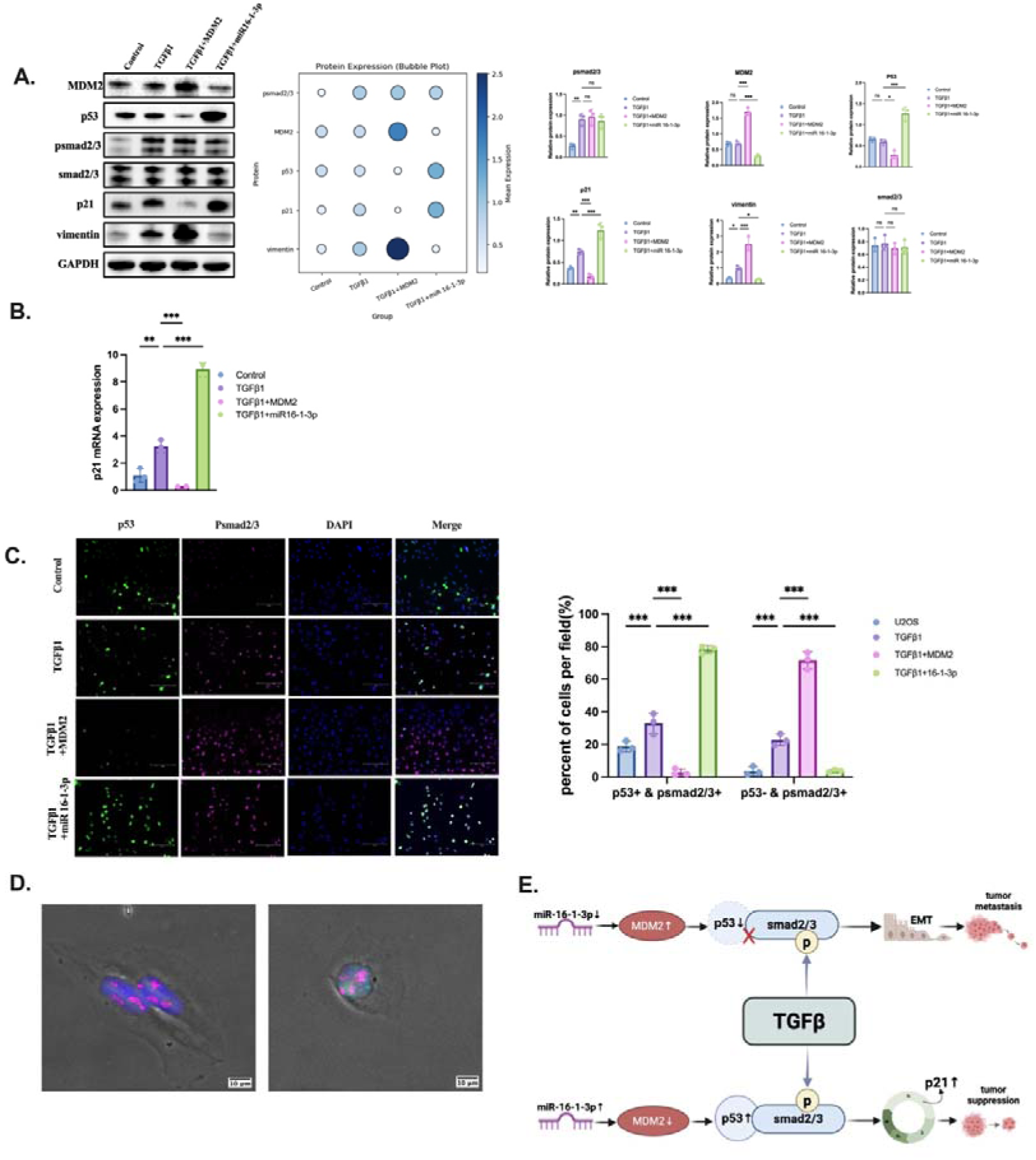
Impact of miR-16-1-3p on the functional output of TGF-β1 signaling. (A) Western blot analysis and bubble plot quantification of U2OS cells transfected with 50□nM miR-16-1-3p mimic, 1□μg MDM2 overexpression plasmid, or control vector. After 24□h, cells were treated with 5□ng/mL TGF-β1 for an additional 24□h. (B) qPCR analysis of p21 mRNA expression in U2OS cells transfected for 24 h with miR-16-1-3p mimic (50 nM) or MDM2 overexpression plasmid, with or without TGF-β1 stimulation (5 ng/mL, 24 h). (C) Dual immunofluorescence staining of pSmad2/3 and p53 (left) and the number of pSmad2/3-positive vs. pSmad2/3–p53 double-positive cells (right) in response to TGF-β1 stimulation. Scale bar= 150 um. (D) Bright-field imaging of pSmad2/3-positive but p53-negative cells (left) and cells co-expressing pSmad2/3 and p53(right), supporting the notion of functional divergence based on signal combination. Scale bar= 200 um. (E) Schematic model: In the presence of functional p53, TGF-β1-induced pSmad2/3 cooperates with p53 to activate p21 expression, leading to growth suppression. However, when p53 is repressed by MDM2, TGF-β1 signaling favors EMT and vimentin upregulation, promoting migration. miR-16-1-3p counters this effect by targeting MDM2, restoring p53 activity, and redirecting TGF-β1 output toward tumor suppression.

Single-cell high content immunofluorescence results further validated this regulatory mechanism. TGF-β1 stimulation significantly raised the number of pSmad2/3-positive cells compared to non-stimulated cells, but the number of p53-positive cells stayed the same. This resulted in an increased proportion of pSmad2/3□/p53□ cells, along with a concurrent rise in pSmad2/3□/p53□ cells. MDM2 overexpression significantly lowered p53 levels, resulting in fewer pSmad2/3□/p53□ cells and more pSmad2/3□/p53□ cells. In contrast, miR-16-1-3p overexpression significantly raised p53 expression, boosting the count of pSmad2/3□/p53□ while significantly decreasing pSmad2/3□p53□ cells (Figure 4C).

Further morphological observations support that the functional output of pSmad signaling depends on the p53 expression background. pSmad2/3□/p53□ cells had a round or epithelial shape, while pSmad2/3□/p53□ tended to be spindle-shaped or mesenchimal-like (Figure 4D).

Together, the results suggest that the intact p53 conducts TGF-β1-activated Smad signaling, leading to anti-proliferative effects through inducing p21. The decrease of p53 by MDM2 modifies TGF-β1-Smad signaling, boosting cell migration and EMT, reflected in rising vimentin and falling p21 levels. The inhibition of MDM2 by miR-16-1-3p leads to the restoration of p53 expression. This restoration, in turn, triggers the TGF-β1-Smad signaling, significantly enhancing growth inhibition and enabling effective tumor suppression (Figure 4E).

### miR-16-1-3p Modulates TGF-**β** Signaling Independently of p53, While MDM2 Requires p53 to Alter TGF-**β** Responses

To explore if p53 and MDM2 influence miR-16-1-3p’s anti-proliferative action, we created a p53 knockout U2OS cell line using CRISPR/Cas9 system, and verified p53 was fully eliminated via Western blot (Figure 5A).Given these conditions, both colony formation and SRB proliferation assays indicated that TGF-β1 lost its growth-inhibitory properties and instead significantly fostered cell proliferation. This suggests that the anti-proliferative phenotype induced by TGF-β1 is dependent on p53’s presence. Even without p53, miR-16-1-3p effectively suppressed colony formation and cell growth, and MDM2 overexpression did not reverse this effect (Figure 5B). These findings indicate that miR-16-1-3p can exert anti-proliferative activity independently of the p53/MDM2 axis.

**Figure 5.**
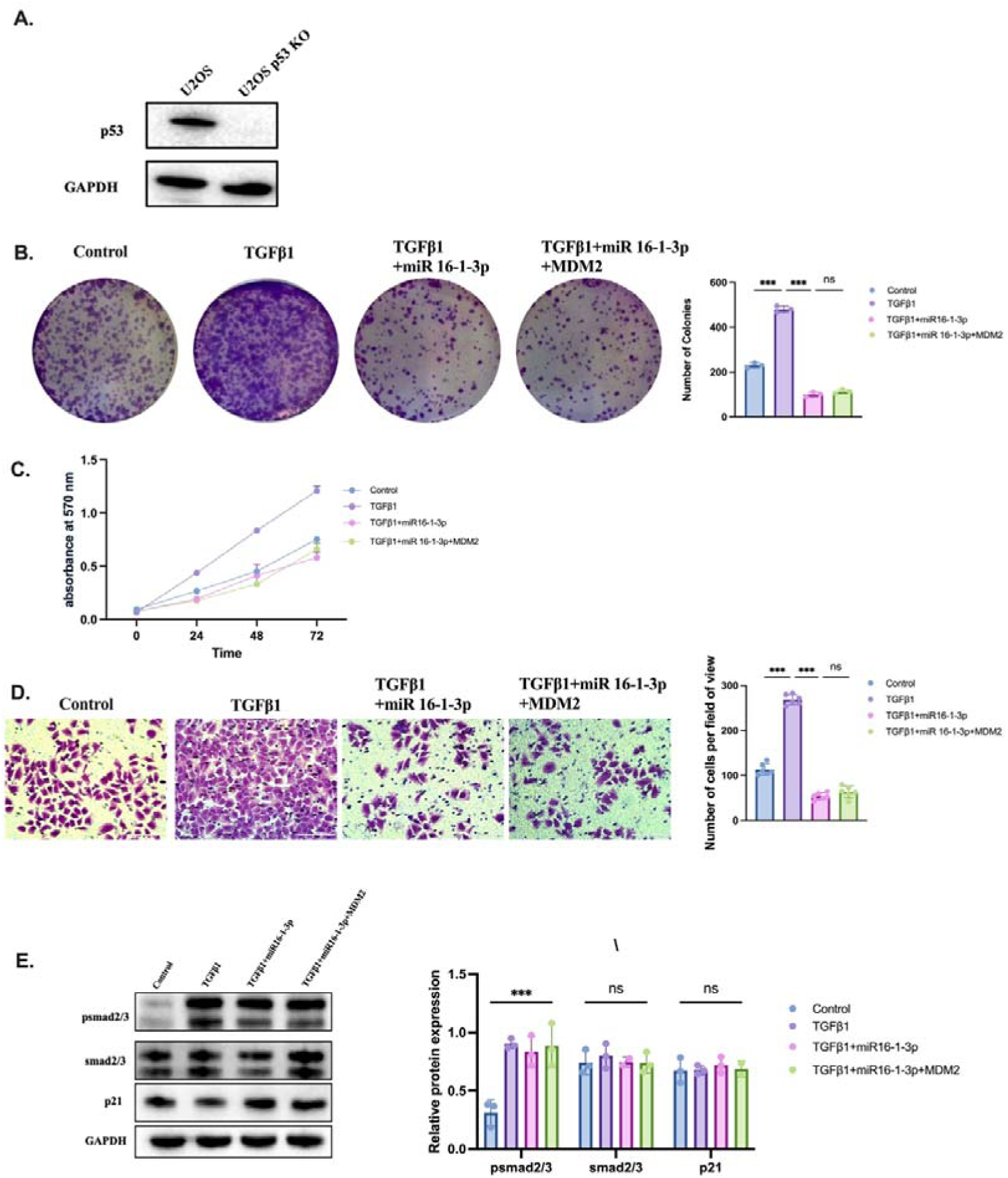
miR-16-1-3p impact on TGF-β1 signaling in p53-knockout U2OS cells. (A) Western blot analysis of p53 expression in wild-type and p53-knockout cells. (B) Colony formation assay: images of crystal violet-stained cell colonies (left) and the number of colonies quantified (right) at 10 days after seeding. (C) SRB proliferation assay: total cell mass (absorbance at 570 nm) was measured at 0, 24, 48, and 72 h after seeding 5000 cells/well in 96-well plate. (D) Transwell assay: Images at 20x magnification display crystal violet-stained cells on the outer membrane of the inner chamber (left). The cell migration rate 24 hours after seeding 100000 cells/well in the upper Boyden chamber of 24-well plate (right). Scale bar = 75 um. (E) Western blot analysis of pSmad2/3, total Smad2/3, and p21expression in cells treated with 5ng/ml TGF-β1 for 24h (left). Quantification of the protein expression (right). GAPDH expression was used for a protein loading control.

TGF-β1 significantly increased cell motility when p53 was absent, indicating that the loss of p53 allows its pro-migratory effects to emerge (Figure 5D). miR-16-1-3p significantly reduced the increased migration, and adding MDM2 did not reverse this effect, reinforcing its tumor-suppressive role in a p53-null environment.

In the absence of p53, TGF-β1 can still promote Smad2/3 phosphorylation, but it fails to enhance p21 protein levels. miR-16-1-3p overexpression, whether alone or with MDM2, had no effect on p21 levels (Figure 5E), implying p53 is essential for TGF-β1’s induction of p21 via the Smad pathway. These results highlight the essential role of p53 as a mediator in the anti-proliferative signaling downstream of TGF-β1.

### miR-16-1-3p redirects TGF-**β**1 signaling to inhibit tumor growth, suppressing metastasis in vivo and chemoresistance in vitro

Changes in TGF-β signaling significantly alter molecule expression, leading to increased cell growth, invasion, EMT, and metastasis in advanced cancers ^26–28^. The chick embryo chorioallantoic membrane (CAM) model improves cancer research by bridging in vitro and in vivo models, offering a cost-effective and accurate way to replicate cancer.^29^ We recently demonstrated that CAM is an efficient model for studying the sensitivity of two breast cancer cell lines to DNA-damaging effect of proton beam irradiation.^29^

Since we found that miR-16-1-3p inhibits both CSCs self-renewal and invasiveness via adjustment of TGF-β1 signaling by targeting MDM2, we explore the in vivo function of miR-16-1-3p. Stably Katushka2S-expressing U2OS cells were transduced with lentivirus constructs to overexpress either miR-16-1-3p or scrambled miRNA (control). Following pretreatment with TGF-β or without it, miR-16-1-3p-expressing cells, along with control cells, were inoculated onto the CAM membranes.

As a result, both TGF-β1 and miR-16-1-3p significantly reduced tumor growth, but the combination of the two had the greatest effect on tumor volume (Figure 6A-B). Immunofluorescence staining provided evidence that TGF-β1 and miR-16-1-3p effectively lowered the number of Ki67-positive cells in tumor nodule tissues, indicating reduced tumor cell growth. As anticipated, TGF-β1 treatment markedly enhanced distant metastasis in the CAM model, while miR-16-1-3p effectively blocked this pro-metastatic effect. The combination group showed significantly reduced metastatic signal compared to TGF-β1 alone, indicating that miR-16-1-3p can counteract the pro-metastatic output of TGF-β1 signaling in vivo (Figure 6C).

**Figure 6.**
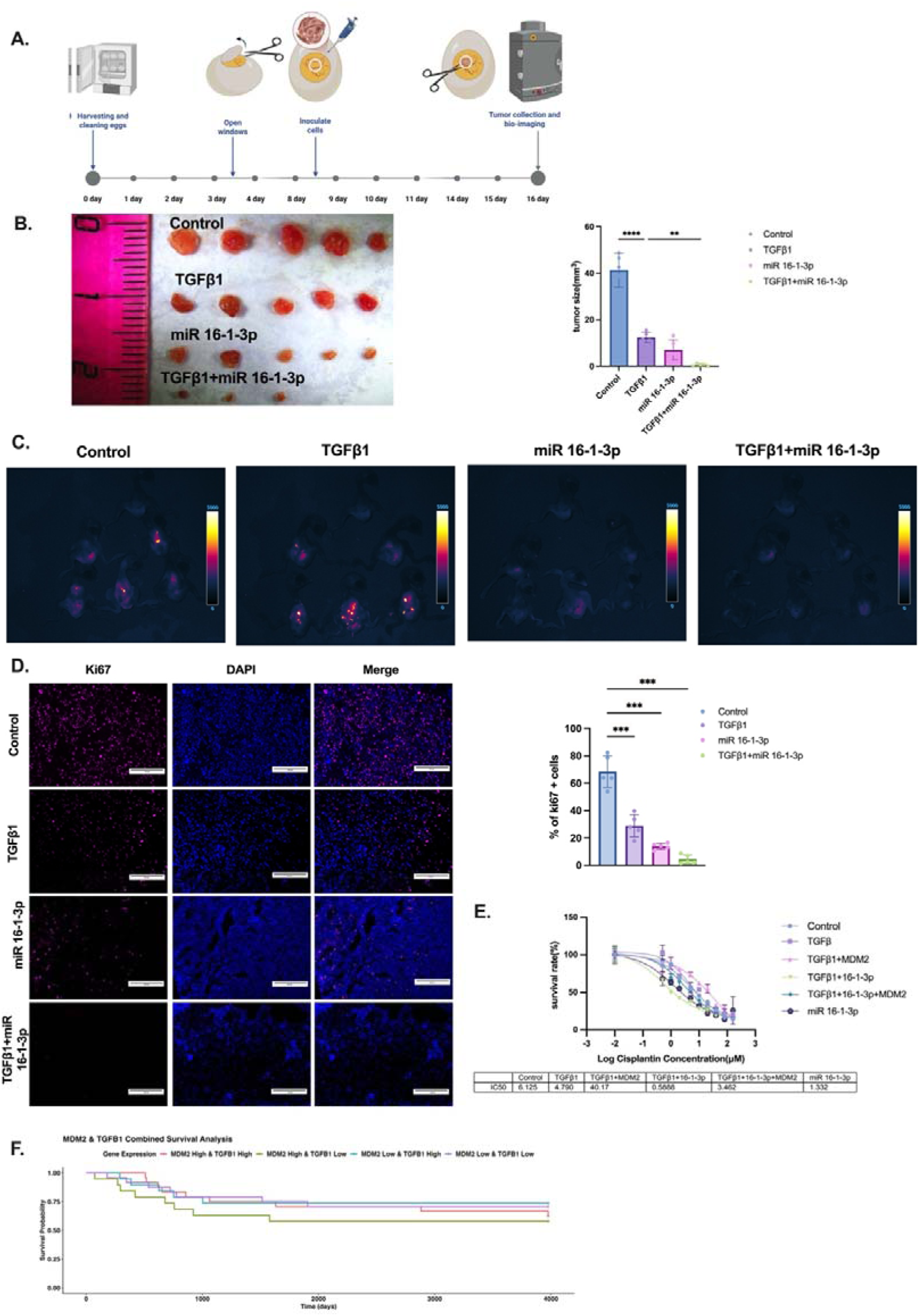
miR-16-1-3p counteracts TGF-β1-mediated effects on metastasis and chemosensitivity of OS cells in vivo. (A) Schematic representation of the in vivo CAM assay. In experiments, stably Katushka2S-expressing U2OS cells (transduced with Katushka2S-expressing lentivirus) were sorted for fluorescent-positive populations. Experimental groups consisted of Katushka2S-positive populations of U2OS cells that were transduced with: PLKO.3G-Scramble (control); miR-16-1-3p (miR-16-1-3p); U2OS cells treated with TGF-β1 (10 ng/10L cells). On the EMD 8 experiment, we implanted 3 × 10L control cells, and cells either lacking (miR-16-1-3p) or containing TGF-β1 (TGF-β1+miR-16-1-3p), onto the CAM to assess tumor growth and metastasis. On EMD 14–16, eggs were imaged and tumors were quantified using LumoTrace® Fluo (Abisense LLC, Russia) imaging system. (B) Representative fluorescent images (left) alongside bright-field macroscopic images (middle) depict the quantification of tumor-associated fluorescence intensities (upper right) and the volumes (lower right) of tumor cell nodules formed on the CAM at EMD 16 post-implantation.(C) Representative fluorescent images of tumor cell metastases formed by Katushka2S-labeled U2OS cells in the chicken embryos at EMD 16 post-implantation. (D) Immunofluorescent analysis of Ki67 expression in the sections of tumor in ovo nodules. Cryosections of tumor nodules were stained with either primary anti-Ki67 (Experimental group) or rat non-specific IgG (Blank) and secondary Anti-Rat IgG H&L, Alexa Fluor®647 antibody (pink color). Pink-purple color represents Ki67 expression, blue represents DAPI-positive staining of nuclei, and last column to the right (Merge) represents overlay images obtained in cyan and blue channels. Amount of Ki67+ cells (Graph on the right) was quantified as % of all cells (DAPI+) using ImageJ software. Scale bar= 150 um (E) SRB assay of dose-dependent proliferation response of U2OS cells to cisplatin (Upper graph). IC50 values (Bottom table) were calculated using built-in algorithms of GraphPad^TM^ software. (F) Kaplan–Meier survival analysis of TARGET-OS patients based on MDM2 and TGFB1 expression. Patients with high TGFB1 and low MDM2 expression showed the best prognosis. Student’s t-test was applied everywhere in order to estimate statistical significance. Error bars represent standard errors of the mean. Two-sided p-value for Student’s t-test: * <0.05 ***<0.001;

Cisplatin sensitivity was assessed using the Sulforhodamine B (SRB) assay using stable miR-overexpressing ( either miR-Scr or miR-16-1-3p) and TGF-β1-treated with or without MDM2 overexpression U2OS cells. Our research demonstrated that TGF-β1 treatment alone did not produce significant changes, while its combination with miR-16-1-3p significantly enhanced cisplatin sensitivity, reducing the IC50 value in U2OS cells by over an order of magnitude in in vitro SRB assays (Figure 6E). The co-transfection of MDM2 partly countered the sensitizing effect, suggesting miR-16-1-3p and TGF-β1 combination boosts chemosensitization of U2OS cells via MDM2 inhibition.

Together, these results indicate that miR-16-1-3p enhances the tumor-suppressive and chemosensitizing effects of TGF-β1 by targeting MDM2, while also mitigating its pro-metastatic output, highlighting its mechanistic importance and potential clinical significance.

To explore the clinical relevance of these findings, we analyzed the prognostic impact of TGFB1 and MDM2 expression levels in the TARGET-OS dataset. Kaplan–Meier survival analysis revealed that patients with high TGFB1 and low MDM2 expression had the most favorable prognosis, whereas low TGFB1 or high MDM2 expression was significantly associated with poor survival (Figure 6F). Together, these results indicate that miR-16-1-3p enhances the tumor-suppressive and chemosensitizing effects of TGF-β1 by targeting MDM2, while also mitigating its pro-metastatic output, highlighting its mechanistic importance and potential clinical significance.

## Discussion

The TGF-β signaling pathway, characterized by its dual ability to suppress proliferation while promoting metastasis under different cellular contexts, has long been recognized as one of the most complex and controversial regulatory modules in cancer biology.^30,31^ Recent studies in gastric cancer have confirmed this functional duality: TGF-β2 activates NDRG1 through a TGFBR/Smad2/3-dependent pathway, enhancing EMT and migration, while simultaneously suppressing proliferation through the same cascade.^32^ Consequently, clinical application of TGF-β inhibitors remains limited due to their inability to selectively block the pro-tumorigenic effects without impairing the tumor-suppressive functions of the pathway. This challenge underscores the urgent need to identify molecular “conductors” that orchestrate the direction of TGF-β signaling output.^3^

At least some of these conductors may exist within the recognized link between TGF-β signaling and p53, crucial for cancer progression.^33^ Studies show that p53 enhances TGF-β signaling by supporting Smad complexes, improving Smad-DNA binding, and regulating tumor-suppressor genes.^16,34^ Functioning as a transcription factor, p53 cooperates with pSmad2/3 to enhance p21 promoter activity, thereby amplifying the growth-inhibitory effect of TGF-β.^16,35,36^ Previous studies have demonstrated that the anti-proliferative activity of TGF-β is dependent on the presence of p53.^3,17,37^ The cyclin-dependent kinase inhibitor p21, a classical Smad2/3 downstream target, is essential for TGF-β-induced cell-cycle arrest. The balance between TGF-β-induced cell-cycle inhibition and EMT activation is largely governed by the status of p53. ^12,16,38^ Collectively, these findings indicate that p53 may functions as a molecular “conductor” determining the phenotypic outcome of TGF-β signaling.

In our p53-wild-type U2OS osteosarcoma model, TGF-β1-Smad activation induced p21 (Fig.3A), resulting in growth inhibition (Fig.1A, B) through cell cycle arrest (Fig.3B). Furthermore, after TGF-β1-Smad activation, U2OS cells changed to a mesenchymal shape, displaying elongated forms and prominent actin stress fibers, as shown by phalloidin staining (Figure 3C). This data together with upregulation of vimentin, featuring epithelial–mesenchymal transition, may explain observed enhanced 2D collective and 3D confined migrations (Fig.1C, D). These data align with findings that activating the TGFβ/Smad signaling pathway causes cytoskeletal alterations and boosts vimentin and myosin levels, enhancing migration in NSCLC cells.^39^ Our data also confirms that TGF-β improves invasiveness by increasing vimentin levels through Smad signaling (Fig.2A). Agents identified through tumorsphere assays have greater potential for in vivo anti-tumor activity than those from general monolayer cultures.^40^ For the first time, we demonstrate that TGFβ/Smad signaling activation also augments invasion, but reduced CSC self-renewal in collagen-embedded 3D tumorsphere assay (Fig.3D) of human osteosarcoma cells. Our research validates and extends prior knowledge on melanoma cells, showing that TGFβ has a major negative effect on stem cell maintenance in various patient-derived cell lines.^41^ Thus, TGF-β1 has been found to inhibit tumor sphere formation, indicating its role in reducing stem cell traits, which is significant in the early progression of tumors. However, TGFβ/Smad signaling activation led to increased CSC invasiveness in OS cell line (Fig.3D).

In p53-knockout U2OS cells, TGF-β1 continued to activate Smad2/3 phosphorylation but failed to induce p21 (Fig. 5E). In mouse colorectal cancer models, p53 gain-of-function mutations or loss of heterozygosity redirect Smad complexes toward the transcription of pro-migratory genes such as Hmga2 and Twist1.^42^ Respectively, p53-knockout U2OS cells had higher growth and clonogenic activity (Fig.5B, C), as well as enhanced 3D confined migration (Fig. 5D), when treated with TGF-β1. These data is in accord to the studies where in p53-deficient or mutant contexts, the anti-proliferative effect of TGF-β was markedly weakened.^38,43,44^ Together, these data suggest that the anti-proliferative vector of TGF-β-Smad signaling is at least partially p53-dependent, whereas the pro-migratory vector of the same signaling is p53-independent. Consequently, ensuring the stabilization of p53 is key to masterfully conducting the complex interactions of TGF-β signaling.

However, the potential of TGF-β in regulating p53 stability still needs of comprehension. The stability of p53 is primarily influenced by the critical processes of ubiquitination and deubiquitination, targeting it to proteasomes. Keeping these in mind, our study focused on MDM2 as a regulator of p53 stability. MDM2, the key E3 ubiquitin ligase for p53, is well known for connecting with p53. This interaction helps ubiquitinate and degrade p53, making MDM2 crucial for inactivating p53 in tumors. Notably, high MDM2 expression in human cells leads to TGF-β1 resistance in various breast cancer cell lines, promoting tumor growth by inhibiting two key tumor suppressors, p53 and Rb.^45^ Curiously, the activation of Smad3/4 regulates the *MDM2* gene, causing an increase in MDM2 protein levels at 48 hours and dramatically increased after 72 hours of TGF-β1 treatment of non-malignant murine mammary epithelial cell line, NMuMg. A rise in MDM2 levels following TGF-β1 treatment coincided with Smad3 activation and the development of the mesenchymal phenotype in NMuMg cells.^17^ This time schedule apparently very well aligns with the dynamics of human U2OS response to TGF-β1 treatment in our collagen-embedded 3D tumorsphere assay, whereby significant increase in invasiveness, albeit reduction of tumorsphere area was observed at 72 hours (Fig.3D). However, no changes in MDM2 or p53 levels, and in cisplatin sensitivity were observed in this OS cell line following TGF-β1 treatment (Fig. 4A and 6E). In contrast, an additional MDM2 overexpression changes how U2OS cells respond to TGF-β1, significantly blocking G1 arrest (Fig.3B) and augmenting tumorsphere area (self-renewal) and invasiveness (peripheral migration) cancer stem cells (Fig.3D) by decreasing p53 and p21 (Fig.4A, B), while enhancing vimentin expression (Fig.3E and 4A) compared to TGF-β1 alone. The same overexpression along with TGF-β1 treatment also dramatically increases 2D collective migration (Fig.1) and reduces the cisplatin sensitivity of U2OS cells by 10-times (Fig.6E). Collectively, the increase of MDM2 protein through Smad3/4 appears inactive in U2OS malignant cells, unlike in non-malignant mouse cells. Nevertheless, constitutive MDM2 protein levels can be instrumental conducting either cytostatic or pro-invasive vectors of TGF-β-Smad signaling in malignancy.

The ultimate clinical marker of a tumor’s ability to migrate and metastasize is the impairment of p53 function. The p53 functions as a transcriptional regulator for the MDM2 gene. However, in advanced stages of metastatic disease, there is a notable increase in MDM2, which usually occurs independently of p53 status. Hence, it seems there are more factors elevating MDM2 protein levels, pointing to a complicated regulatory network.^17^ This prompted us to conduct further research to fully elucidate these alternative pathways. Understanding these mechanisms could provide deeper insights into the regulation of MDM2 and its implications in cellular malignant processes. Overall, this highlights the need to explore additional factors that may influence MDM2’s expression in context of TGF-β signaling.

Recent studies highlight non-coding RNAs (ncRNA) as important regulators of cancer pathways, leading to targeted treatment strategies.^46^ The role of ncRNAs in modulating TGF-beta 1 signaling is complex and context-dependent.^47^ While some ncRNAs facilitate metastasis via EMT, others help suppress tumors by regulating TGF-beta 1’s growth-inhibiting properties. Attempting to block TGF-β without knowing its dual effects could compromise its tumor-suppressing functions. Hence, finding the molecular switches that regulate TGF-β signaling is vital for minimizing its cancer-promoting effects while keeping its benefits against tumors.

An exonic ncRNA, circNUDT21 acted as a miR-16-1-3p sponge, and increasing MDM2 helps downregulate p53, promoting bladder cancer.^48^ Our recent bio-informatic study ^49^ and present data (Fig.2A, B) confirmed that MDM2 is one of direct targets of miR-16-1-3p in U2OS cells. It is essential to highlight that although miR-16-1-3p overexpression by itself markedly reduces proliferative and clonogenic potential of U2OS cells, when paired with TGF-β treatment, it significantly increase arrest cells in G1 phase (Fig. 3B) and nearly extinguishing the growth capability of these cells (Fig.2C, D). Furthermore, miR-16-1-3p overexpression inhibited TGF-induced actin remodeling and EMT featuring, significantly decreasing vimentin and increasing p21 levels shown in Fig. 3C and Fig. 3E. Importantly, TGF-β enhances both 2D and 3D migration, but miR-16-1-3p overexpression, alone or with TGF-β, strongly counteracts its migration effects (Fig. 2E, F). Conversely, all effects of miR-16-1-3p were significantly remedied by additional MDM2 overexpression (Fig.2C-F and Fig. 3B-E), confirming the existence of TGF-β-MDM2 regulatory network.

Importantly, miR-16-1-3p restored p53 stability by targeting MDM2, redirecting TGF-β-Smad signaling toward p21 activation (Fig. 4A-C) and proliferation inhibition while attenuating its EMT-promoting capacity (Fig. 4D). In osteosarcoma cells lacking functional p53, treatment with miR-16-1-3p had similar effects even in the presence of additional MDM2 expression (Fig. 5).

Recent studies reveal that non-coding RNAs significantly contribute to oxaliplatin resistance of colorectal cancer cells via transcriptional, post-transcriptional, and epigenetic mechanisms.^50^ In a groundbreaking finding, we demonstrated that the simultaneous administration of TGF-β and miR-16-1-3p dramatically increases the sensitivity of U2OS cells to cisplatin, exceeding that of TGF-β therapy alone by more than an order of magnitude (Fig. 6E). Changes in TGF-β signaling significantly alter molecule expression, leading to increased invasion, EMT, metastasis and chemoresistance in advanced cancers.^27,28,51,52^ However, it remains to be determined if this signaling is functional in the CAM in vivo model. For the very first time, administering TGF-β and miR-16-1-3p together significantly reduces the tumor nodule volume and Ki67 expression (Fig. 6B), while effectively eradicates metastases (Fig. 6C) in the CAM in vivo model. These data corroborate our observed anti-migratory in vitro effects. Unlike small-molecule MDM2 inhibitors, miR-16-1-3p acts post-transcriptionally, avoiding their dose-dependent toxicity and resistance ^53^ while concurrently modulating both p53 and TGF-β functional outcomes.

Noteworthy, TGF-β reveals its versatility by signaling through non-Smad pathways as well.^6^ TGF-β activates mTOR via the PI3K/AKT pathway, enhancing ubiquitin carboxyl-terminal hydrolase 15 (USP15) translation. USP15 then binds to and stabilizes p53 by removing its ubiquitin tags. USP15 caused the MDM2/p53 complex to break apart, creating a new MDM2/USP15 complex.^6^ Thus, the PI3K/AKT pathway may aid TGF-β signaling in enhancing p53 stability by boosting USP15 translation, which could clarify its role in inhibiting early cancer in cells with normal p53.

## Conclusion

Understanding how different ncRNAs interact with the TGF-beta 1 pathway is essential for creating effective cancer therapies aimed at either enhancing tumor suppression or preventing metastasis. More importantly, it’s unclear what upstream factors affect this switch, how they change during disease progression, and if they can be targeted for treatment. This fundamental gap in knowledge has hindered the development of rational TGF-β-based therapeutic strategies. Our study establishes that the functional polarity of TGF-β signaling depends on p53 status and identifies the miR-16-1-3p–MDM2–p53 axis as a key conductor orchestrating cytostatic and pro-invasive vectors of TGF-β-Smad signaling in malignancy. We demonstrate that restoring p53 stability via miR-16-1-3p can selectively preserve the tumor-suppressive and anti-proliferative functions of TGF-β while limiting its pro-metastatic effects. These findings provide mechanistic insight into the long-standing paradox of TGF-β signaling and offer a promising therapeutic strategy for precision modulation of TGF-β activity in osteosarcoma and possibly other solid tumors.

## Limitations of current study

This study systematically elucidated the critical role of the miR-16-1-3p–MDM2–p53 axis in regulating the functional output of TGF-β signaling and demonstrated, through in vitro and in vivo experiments, its biological significance in balancing the anti-proliferative and pro-migratory effects in osteosarcoma cells. However, several limitations should be acknowledged and addressed in future work:

First, our results indicate that the downstream effects of TGF-β-induced pSmad2/3 are at least partially dependent on the presence of intact p53. However, it is still unclear if miR-16-1-3p’s reduction of MDM2, meant to potentially slow tumor growth by targeting MDM2–Rb axis, also directly affects TGF-β1 sensitivity in cancers.

Second, the clinical data analyzed in this study were derived primarily from the public TARGET-OS database, which has a relatively limited sample size and incomplete records of chemotherapy response and long-term follow-up. This limitation constrains the comprehensive evaluation of the prognostic value of the miR-16-1-3p–MDM2–p53 axis across different clinical subtypes or treatment stages. Future studies incorporating larger patient cohorts and histological validation are warranted to confirm the clinical relevance and translational potential of this signaling pathway.

In summary, this study proposes a novel mechanistic hypothesis and establishes a solid experimental framework, but further molecular and clinical investigations are required to advance from mechanistic discovery toward the development of precise, target-based therapeutic strategies.

## Materials and Methods

### Cell culture

The human osteosarcoma cell line U2OS and the human embryonic kidney cell line HEK293T were obtained from the Shared Research Facility “Vertebrate cell culture collection” at the Institute of Cytology, Russian Academy of Sciences (Saint Petersburg, Russia). Cells were maintained in high-glucose Dulbecco’s Modified Eagle’s Medium (DMEM, Gibco) supplemented with 10% fetal bovine serum (FBS, Gibco), 100 U/mL penicillin, and 100 μg/mL streptomycin. Cultures were incubated at 37 °C in a humidified atmosphere with 5% CO□. All cell lines were routinely tested for mycoplasma contamination and only cells in logarithmic growth phase were used for experiments.

### Plasmid Construction

For the construction of the MDM2 overexpression plasmid, standard molecular cloning procedures were performed. The pcDNA3.1(+) vector was double-digested with the restriction enzymes BamHI and XhoI (Thermo Fisher Scientific). The human MDM2 coding sequence was amplified by PCR and digested with the same enzymes to generate compatible cohesive ends. The digested vector and insert were ligated using T4 DNA ligase. The ligation products were transformed into competent E. coli cells by heat shock and plated on antibiotic-containing agar plates. Positive colonies were screened by colony PCR, and the presence, orientation, and integrity of the MDM2 insert were confirmed by DNA sequencing. Plasmid DNA was subsequently isolated from bacterial cultures using the GeneJET Plasmid Miniprep Kit (Thermo Fisher Scientific) following the manufacturer’s instructions.

Similarly, a lentiviral plasmid containing the miR-16-1* and shScr sequences was constructed. The lentiviral vector pLKO.3G was kindly provided by Christophe Benoist and Diane Mathis (Addgene plasmid #14748; http://n2t.net/addgene:14748; RRID: Addgene_14748). Lentiviral constructs pLKO.3G–miR-16-1* and pLKO.3G–Scr miR, containing the EGFP reporter gene for visualization and designed for overexpression of miR-16-1* and Scr miR, respectively, were generated as follows.

Duplexes of miR-16-1* were synthesized by annealing complementary oligonucleotides hsa-miR-16-1*-F and miR-16-1*-R (see Supplementary Table□S1). The resulting duplexes were cloned into the Acc36I and EcoRI restriction sites of the pLKO.3G vector, yielding the corresponding pLKO.3G–miR-16-1* constructs. The accuracy of the cloned lentiviral constructs was verified by PCR using primers pLKO-Dir and pLKO-Rev, followed by Sanger sequencing with primers PLKO.1 5′ and pLKO-Rev to confirm insert orientation and sequence integrity. The sequence of the scrambled miRNA was previously published. ^54^

### CRISPR/Cas9-mediated TP53 knockout

To generate TP53-knockout U2OS cells, an sgRNA targeting an early exon of the human TP53 gene was designed using the Synthego Knockout Guide Design Tool. The selected sgRNA sequence was GCATGGGCGGCATGAACCGG. The lentiCRISPR v2 puro plasmid (Addgene #98290, a gift from Brett Stringer) was used as the backbone vector. Oligonucleotides containing the sgRNA sequence (forward: 5′-CACCGGCATGGGCGGCATGAACCGG-3′; reverse: 5′-AAACCCGGTTCATGCCGCCCATGC-3′) were annealed and ligated into the BsmBI-digested vector. The cloned product was verified by Sanger sequencing using the hU6-F primer (5′-GAGGGCCTATTTCCCATGATT-3′), confirming correct sgRNA insertion downstream of the U6 promoter.

The verified construct was co-transfected with pLP1, pLP2, and pVSVG packaging plasmids into HEK293T cells to produce lentiviral particles. Viral supernatants were collected, filtered, and used to infect U2OS cells in the presence of 8□µg/mL polybrene. Forty-eight hours post-infection, cells were selected with 1□µg/mL puromycin for approximately 7□days to establish a stable population. The knockout was validated by Sanger sequencing of the target region and confirmed functionally by Western blot, showing complete loss of P53 protein expression.

### Transfection, Lentiviral Transduction and Treatment

For MDM2 overexpression, U2OS cells were transfected with an MDM2 overexpression plasmid using Lipofectamine™ 3000 (Invitrogen, USA) according to the manufacturer’s protocol. Plasmid DNA was mixed with P3000 reagent and Lipofectamine 3000 in Opti-MEM and added to cells at 60–70% confluency. After 24 hours, cells were collected for subsequent assays. The efficiency of MDM2 overexpression was confirmed by qRT-PCR, as shown in Supplementary Figure 1A. For in vitro experiments requiring transient miRNA manipulation, U2OS cells were transfected with either miR-16-1-3p mimic (50 nM) or Scramble mimic (Syntol LLC, Russia, Moscow) using Lipofectamine™ 3000. After 24 hours of transfection, cells were harvested or reseeded for downstream assays. Transfection efficiency of miR-16-1-3p mimic was verified by qRT-PCR, and the results are shown in Supplementary Figure 1B.

For generating fluorescently traceable cell lines for CAM xenograft assays, U2OS cells were first infected with lentivirus encoding the red fluorescent protein Katushka2S. After 72 hours, Katushka2S-positive cells were isolated by fluorescence-activated cell sorting (BIO-RAD S3e). The purified Katushka2S□ population was subsequently infected with GFP-labeled lentiviruses expressing either miR-16-1-3p (PLKO.3G-miR-16-1-3p) or Scramble control (PLKO.3G-miR-Scr). Following another 72-hour incubation, GFP-positive cells were sorted to obtain double-positive (Katushka2S□/GFP□) stable lines. The efficiency of stable miR-16-1-3p overexpression in these dual-labeled cells was confirmed by qRT-PCR, as shown in Supplementary Figure 1C.

In experiments involving MDM2 overexpression, cells in the control and comparison groups were transfected with an empty vector plasmid to ensure consistency. In all experiments involving miR-16-1-3p, a scrambled miRNA was used in the control group to account for nonspecific effects. For TGF-β1 treatment, recombinant human TGF-β1 (Novoprotein, China) was diluted in complete medium to a final concentration of 5 ng/mL and added directly to the culture medium as part of the experimental treatment.

### Colony formation assay

The colony formation ability was evaluated using a plate colony formation assay. Cells were seeded in 6-well plates at a density of 500 cells per well and cultured in complete medium containing 10% FBS for 10–14 days until visible colonies were formed. The medium was removed, and cells were gently washed twice with PBS, fixed with 4% paraformaldehyde for 15 min, and stained with 0.1% crystal violet for 30 min. Excess dye was washed off, and the plates were air-dried. Colonies were photographed and counted, and the colony formation efficiency was calculated as number of colonies.

### SRB proliferation assay

Cell proliferation was assessed using the Sulforhodamine B (SRB) assay. Cells were seeded in 96-well plates at a density of 3000–5000 cells per well. At 0, 24, 48, and 72 h, cells were fixed by adding trichloroacetic acid (TCA) to a final concentration of 10% and incubating at 4 °C for 1 h. Plates were washed with distilled water and air-dried. Cells were then stained with 0.4% SRB solution for 30 min at room temperature. Excess dye was removed by washing with 1% acetic acid, and plates were air-dried. Bound dye was solubilized with 10 mM Tris-HCl (pH 10.5), and absorbance was measured at 510 nm using a microplate reader. Cell proliferation was quantified and plotted as growth curves based on absorbance values at each time point.

### Transwell migration assay

3D confined cell migration capacity was assessed using 24-well Transwell chambers with 8 μm pore polycarbonate membranes (SPLInsert™ Hanging, PET membrane, Cat no.37124). Serum-starved cells (1 × 10^5^ per well) were suspended in serum-free medium and seeded into the upper chamber, while the lower chamber was filled with complete medium containing 10% FBS as a chemoattractant. After incubation for 24 h at 37 °C in a humidified atmosphere with 5% CO□, non-invading cells on the upper surface were removed with a cotton swab. The membranes were fixed with 4% paraformaldehyde and stained with 0.1% crystal violet. Invaded cells were counted under a light microscope (Leica DFC 7000 Т) in three randomly selected fields, and the average was used for statistical analysis.

### Wound healing assay

2D collective cell migration was assessed by wound healing assay. Cells were seeded into 6-well plates and grown to approximately 90% confluence. A straight scratch was made across the monolayer using a sterile 200 μl pipette tip, and detached cells were removed by washing with PBS. Cells were then incubated in serum-free medium. Images of the wound area were captured at 0 h and 24 h by the EVOS™ M5000 Imaging System, and the scratch area was quantified using ImageJ software (National Institute of Health, USA). The migration rate was calculated as: (Initial wound area – Wound area at 24 h) / Initial wound area × 100%.

### Dual-luciferase test

To validate the ability of hsa-miR-16-1 to interact with the predicted target site within the MDM2 mRNA, a dual-luciferase reporter assay was performed using the pmiR-Glo vector system, specifically designed for studying gene expression regulation mediated by microRNA binding to the 3′ untranslated regions (3′-UTRs) of mRNAs. The predicted miR-16-1 binding site was amplified by polymerase chain reaction (PCR) and subsequently cloned into the multiple cloning site of the pmiR-Glo plasmid, located downstream of the firefly luciferase gene, between the NheI (Thermo Fisher) and XhoI (Thermo Fisher) restriction sites. Following restriction digestion of both the vector DNA and the amplified fragment with the corresponding endonucleases, ligation was carried out using T4 DNA ligase (Thermo Fisher).Transformed *E.coli* colonies were screened for the presence of the insert by PCR analysis, and positive clones were further verified by DNA sequencing. Plasmid DNA was then purified using the GeneJET Plasmid Miniprep Kit (Thermo Fisher Scientific). For the functional assay, 300 ng of the reporter plasmid were co-transfected with concentration of 50nM hsa-miR-16-1 mimics using Lipofectamine 3000 (Thermo Fisher Scientific), according to the manufacturer’s instructions. After 24 hours, luciferase activity was measured using the Dual-Lumi™ Luciferase Reporter Gene Assay Kit (Servicebio, China)and quantified with a ClarioStar spectrophotometer (BMG Labtech). All measurements were performed in triplicate, and firefly luciferase activity was normalized to Renilla luciferase activity to control for transfection efficiency.

### Cisplatin sensitivity assay

Cisplatin sensitivity was assessed using the Sulforhodamine B (SRB) assay. U2OS cells were first transfected for 24 h with either Scramble mimic or miR-16-1-3p mimic (50 nM), with or without co-transfection of the MDM2 overexpression plasmid, and treated with TGF-β1 (5 ng/mL) according to the experimental design. After pretreatment, cells were seeded into 96-well plates at a density of 10,000 cells per well and allowed to adhere overnight. Cisplatin was then added at concentrations ranging from 0.01 to 100 μM (logarithmic serial dilutions). After 72 h of treatment, cells were fixed with 10% trichloroacetic acid (TCA) at 4 °C for 1 h, washed, and air-dried. Cells were stained with 0.4% SRB solution for 30 min at room temperature, washed with 1% acetic acid, and dried. The bound dye was solubilized with 10 mM Tris-HCl (pH 10.5), and absorbance was measured at 510 nm using a microplate reader. Cell viability was normalized to untreated controls (100%), and IC50 values were determined by nonlinear regression analysis of dose–response curves using GraphPad Prism 10.0.

### 3D Tumorsphere assay

We created a 3D tumor model with fluorescence and studied it using stable Katushka2S/miR-overexpressing (MiR-Scr/MiR-16-1-3p) U2OS cells, treated or untreated with TGF-β1, and with or without MDM2 overexpression. These cells were seeded in agarose-embedded 96-well plates to monitor tumor spheroid growth and migration continuously. First, 1.5% sterile agarose was added to 96-well plates as a bottom layer. After solidification, 20,000 cells were seeded per well and centrifuged to promote aggregation into compact spheroids. The spheroids were cultured in agarose for 48 hours, and fluorescence microscopy was used to image and quantify the spheroid area, reflecting 3D proliferation capacity.^55^ Next, spheroids were transferred into a 3D collagen matrix composed of 1□mg/mL Rat Tail Type I collagen (BD Biosciences) and overlaid with serum-free medium for an additional 24 hours.^56^ During this phase, the expansion of fluorescence signals from the spheroid edges was recorded to assess invasion and migration through the extracellular matrix. Fluorescent areas were quantified using ImageJ software, with the increase in signal area representing tumor spheroid proliferation and invasive outgrowth. Fluorescence areas were quantified using ImageJ software. Proliferation and migration rates were calculated as:

(Final fluorescence area − Initial fluorescence area) / Initial fluorescence area.

### Cell cycle analysis

Cells subjected to different treatments were harvested and washed twice with PBS. The cells were then fixed overnight at 4 °C in 70% pre-chilled ethanol. After fixation, the cells were washed with PBS and stained with a solution containing RNase A (100 μg/ml) and propidium iodide (PI, 50 μg/ml), followed by incubation at 37 °C for 30 min in the dark. The samples were analyzed by flow cytometry (BD Biosciences, USA). DNA content and cell cycle distribution (G0/G1, S, G2/M phases) were quantified using FlowJo software (Tree Star, USA).

### CAM Tumor Formation and Metastasis in vivo Assay

We implemented our well-established and thoroughly documented protocol ^57^. Briefly, fertilized specific pathogen-free (SPF) eggs were obtained from a certified local hatchery (Trade house Ptichnoe, Ltd., https://ptichnoe-td.ru). Eggs were wiped with sterile water and non-woven paper towels, then incubated horizontally at 37 °C and 70% relative humidity with automatic rotation, designated as embryonic day 0 (EMD 0). On EMD 3, the air sac and major blood vessels were visualized using an egg candler, and approximately 3 mL of albumen was withdrawn to lower the chorioallantoic membrane (CAM). A circular window of 2 cm in diameter was opened in the shell using a mini rotary saw, sealed with 3M semipermeable film, and returned to the incubator with rotation disabled.

On EMD 8, tumor cells were seeded onto a vascular-rich region of the CAM within a polytetrafluoroethylene O-ring (inner diameter 6 mm, outer diameter 9 mm). A 25 μL suspension containing 3 × 10□ cells was carefully applied. All cells used in this assay were stably transduced with Katushka-expressing lentivirus and sorted for fluorescent-positive populations. Experimental groups consisted of control U2OS cells stably transduced with PLKO.3G-Scramble; U2OS cells overexpressing miR-16-1-3p (PLKO.3G-miR-16-1*); U2OS cells treated with TGF-β1, where an equivalent concentration of TGF-β1 (10 ng per 10□ cells) was premixed with 20 μL of the cell suspension immediately before inoculation; and a combined treatment group in which miR-16-1-3p–overexpressing cells were suspended together with the equivalent TGF-β1 concentration. All suspensions were prepared in serum-free medium mixed 1:1 (v/v) with Matrigel (Corning, 356234) to enhance tumor take rate.

After implantation, the windows were resealed and eggs were returned to the incubator. On EMD 16, tumors and embryos were harvested. Primary tumors were imaged using a Leica M60 stereomicroscope, and tumor volume was calculated using the ellipsoid formula:

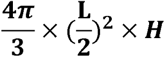

where L represents the maximal tumor diameter and H the perpendicular height. For metastasis analysis, whole embryos were imaged with an in vivo imaging system to detect Katushka fluorescence, and metastatic foci were quantified based on the distribution of distant fluorescent signals.^58^

According to international guidelines, chicken embryos are not considered live animals before day 17 of incubation; therefore, all experiments were terminated at EMD 16 and were exempt from IACUC approval.

### RNA isolation and RNA detection

Total RNA was extracted using TRIzol reagent and quantified for concentration and purity using a NanoDrop spectrophotometer (Thermo Fisher Scientific). For mRNA quantification, one-step SYBR Green RT-qPCR was performed using the BioMaster RT-qPCR SYBR Blue (2×) kit (Biolabmix, RM04-400), with GAPDH serving as the internal control. Specific primers were used to detect MDM2, p21, and GAPDH (primer sequences are provided in Supplementary Table□S2).

For microRNA quantification, reverse transcription was performed using the High-Capacity cDNA Reverse Transcription Kit (Thermo Fisher Scientific, Cat. No.□4368814) with stem-loop primers specific to each microRNA. The expression level of hsa-miR□16□1□3p was quantified using a TaqMan probe-based qPCR assay, employing a microRNA-specific forward primer, a universal reverse primer, and the corresponding TaqMan probe. U44 small nuclear RNA was used as the internal control (all primer and probe sequences are listed in Supplementary Table□S2).

qPCR reactions were performed on an Applied Biosystems QuantStudio□5 Real-Time PCR System under standard cycling conditions (95□°C for□10□min, followed by□40□cycles of□95□°C for□15□s and□60□°C for□1□min). For data normalization, mRNA levels were standardized to GAPDH, and microRNA expression was normalized to U44 small nuclear RNA. Relative expression levels were calculated using the□2^–ΔΔCt□method. All reactions were performed in three independent biological replicates to ensure statistical reliability.

### Western blot analysis

Cells were lysed on ice using RIPA buffer supplemented with protease inhibitors (1:100 dilution), and lysates were centrifuged at 12,000 rpm for 15 min at 4 °C to collect the supernatants. Protein concentrations were determined using a BCA Protein Assay Kit (Thermo Fisher Scientific). Equal amounts of protein were separated by SDS–PAGE and transferred to PVDF membranes, which were blocked with 5% BCA for 1 h at room temperature. Membranes were incubated overnight at 4 °C with the following primary antibodies: MDM2 (1:1000, ab259265), p53 (1:1000, ab32509), p21 (1:1000, ab109520), Vimentin (1:1000, Elabscience), Smad2/3 (1:1000, Thermo Fisher Scientific,PA5-36125), phospho-Smad2/3 (1:1000, Thermo Fisher Scientific, PA5-110155), and GAPDH (1:3000, ZB15004-HRP). After three washes with TBST, membranes were incubated with HRP-conjugated secondary antibodies (1:2000, Affinity S0001) for 60 min at room temperature, followed by three additional TBST washes. Protein bands were visualized using enhanced chemiluminescence (ECL, Bio-Rad, 1705061) reagents, and images were acquired with a ChemiDoc XRS+ system (Bio-Rad). Band intensities were quantified using ImageJ software and normalized to GAPDH expression.

### Immunofluorescence staining

For in vitro staining, U2OS cells were seeded on sterile glass coverslips, fixed with 4% paraformaldehyde for 15 min, and permeabilized with 0.2% Triton X-100 for 10 min. Cells were blocked with 5% bovine serum albumin (BSA) for 1 h at room temperature, then incubated overnight at 4 °C with primary antibodies: anti-p53 (1:200, ab237976) and anti-phospho-Smad2/3 (1:200, Thermo Fisher Scientific, PA5-110155). After washing with PBS, Alexa Fluor 488-conjugated anti-mouse(1:500, ab150113) and Alexa Fluor 647-conjugated anti-rabbit secondary (1:500, ab150079) antibodies were applied for 1 h at room temperature in the dark. Nuclei were counterstained with DAPI, and slides were mounted with anti-fade medium. Images were acquired using the EVOS™ M5000 Imaging System.

For in vivo staining, CAM tumors harvested at embryonic day 16 (EMD 16) were fixed in 4% paraformaldehyde, cryoprotected in 30% sucrose, embedded in OCT compound, and sectioned at 8 µm. Frozen sections were washed with PBS, permeabilized with 0.2% Triton X-100 for 10 min, and blocked with 5% BSA for 1 h. Sections were then incubated overnight at 4 °C with rabbit anti-Ki67 (1:200, Abcam), followed by incubation with Alexa Fluor 647-conjugated anti-rabbit secondary antibody (1:500, Invitrogen) for 1 h at room temperature in the dark. Nuclei were counterstained with DAPI, and fluorescence images were captured using the EVOS™ M5000 Imaging System. The proportion of Ki67-positive cells was quantified using ImageJ software.

### Bioinformatic analysis

Transcriptomic and clinical follow-up data of osteosarcoma patients were obtained from the TARGET-OS database (https://ocg.cancer.gov/programs/target), including 88 samples in total. Gene expression matrices were normalized using log2(FPKM + 1) transformation. Based on TGFB1 and MDM2 expression levels, samples were dichotomized into “high” and “low” groups using the median expression value as the cutoff. For combined analysis, patients were categorized into four subgroups: TGFB1 high/MDM2 low, TGFB1 high/MDM2 high, TGFB1 low/MDM2 high, and TGFB1 low/MDM2 low.

Kaplan–Meier survival curves were generated to assess overall survival differences among the four groups, and significance was determined by the log-rank test. Statistical analyses were performed in R software (version 4.3.0) using the “survival” (v3.5-7) and “survminer” (v0.4.9) packages.

### Statistical Analysis

Statistical analysis was performed using GraphPad Prism version 10 (GraphPad Software, San Diego, USA). All experiments were independently repeated at least three times, and data are presented as mean ± standard deviation (SD). For comparisons between two groups, unpaired two-tailed Student’s t-tests were used. For comparisons involving three or more groups, one-way analysis of variance (ANOVA). A p-value of less than 0.05 was considered statistically significant. The levels of significance were denoted as follows in the figures: *P < 0.05, **P < 0.01 and ***P < 0.001, while “n.s.” indicates no significant difference.

## Acknowledgements

This work acknowledge the financial support from the Russian Science Foundation (project No. 23-14-00220).

## Authorship contributions

**Wenyu Xue**: Writing – review & editing, Writing – original draft, Visualization, Validation, Methodology, Investigation, Formal analysis, Data curation, Conceptualization. **Yuzhe Wang**: Writing – review & editing, Writing – original draft, Validation, Methodology, Formal analysis, Data curation, Conceptualization. **A.V. Smirnova**: Writing – review & editing, Validation, Methodology, Investigation, Data curation. **P.A. Malakhov**: Writing – review & editing, Validation, Methodology, Investigation, Data curation. **Margarita Pustovalova**: Writing – review & editing, Visualization, Validation, Supervision, Conceptualization. **Denis V Kuzmin**: Writing – review & editing, Visualization, Supervision, Resources, Project administration, Funding acquisition. **Sergey Leonov**: Conceptualization, Writing – review & editing, Writing – original draft, Data curation, Visualization, Supervision, Resources, Project administration.

## Data availability

The datasets used and/or analysed during the current study available from the corresponding author on reasonable request.

The bio-imaging data that support the findings of this study are available from Abisense LLC but restrictions apply to the availability of these data, which were used under license for the current study, and so are not publicly available. Data are however available from the authors upon reasonable request and with permission of Abisense LLC.

