## Supplemental Table 1 and 2 Supplemental Figure 1 for "The Small Non-coding RNA miR-16-1-3p Hampers Cancer Stem Cell Self-renewal and Invasiveness, Boosting Chemosensitivity by Adjusting TGF-β1 Signaling via MDM2/p53 Axis in Human Osteosarcoma"

**Supplementary Table 1.Oligonucleotides used for cloning miR-16-1-3p and scrambled control sequences into the PLKO.3G lentiviral vector.**

| Oligonucleotide Name | Oligonucleotide Sequence |
|---|---|
| hsa-miR-16-1*-F | accggCCAGTATTAACTGTGCTGCTGActcgagTCAGCAGCACAGTTAATACTGGtttttg |
| hsa-miR-16-1*-R | aattcaaaaaCCAGTATTAACTGTGCTGCTGActcgagTCAGCAGCACAGTTAATACTGGc |
| shScrambled-F | aCCGGT*CCTAAGGTTAAGTCGCCCTCG*CTCGAG*CGAGGGCGACTTAACCTTAGG*TTTTTG |
| shScrambled-R | AATTCAAAAA*CCTAAGGTTAAGTCGCCCTCG*CTCGAG*CGAGGGCGACTTAACCTTAGG*Ac |
| PLKO-Dir | tgtggaaaggacgaaacacc |
| PLKO-Rev | Tctttcccctgcactgtacc |

**Supplementary Table 2.Primer sequences used for quantitative real-time PCR (qRT-PCR) analysis**

| Name | Primer |
|---|---|
| Human U44 RT Primer | GGTCGTATGCAAAGCAGGGTCCGAGGTATCCATCGCACGCATCGCACTGCATACGCAGTCAGTTAG |
| hsa-miR-16-1-3p RT Primer | GGTCGTATGCAAAGCAGGGTCCGAGGTATCCATCGCACGCATCGCACTGCATACGACCTCAGCAGC |
| Human U44 Dir Primer | TGCTGACTGAACATGAAGGTC |
| hsa-miR-16-1-3p Dir Primer | TCGGCGCCAGTATTAACTGT |
| Universal Rev Primer | GCAAAGCAGGGTCCGAGG |
| Taq man probe | FAM 5′ TCCATCGCACGCATCGCACT 3′ BHQ-1 |
| MDM2-F | ACTGTGTATCAGGCAGGGGA |
| MDM2-R | AAGCCCTCTTCAGCTTGTGT |
| hCDKN1A-F | GCCGAAGTCAGTTCCTTGTG |
| hCDKN1A-R | TCGAAGTTCCATCGCTCACG |
| GAPDH-F | ACATCGCTCAGACACCATG |
| GAPDH-R | TGTAGTTGAGGTCAATGAAGGG |

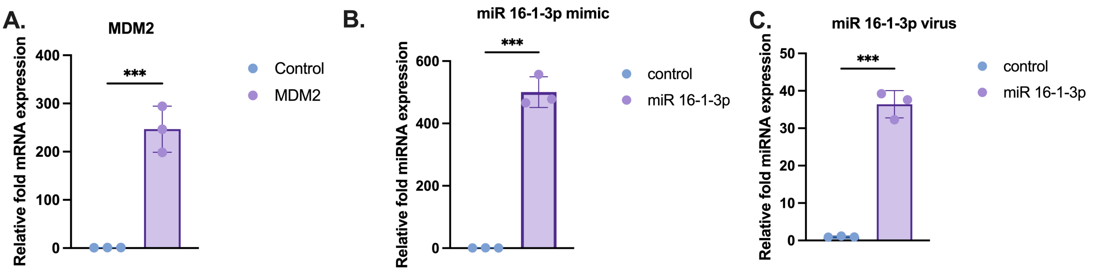


**Supplementary Figure 1. Validation of MDM2 and miR-16-1-3p overexpression by qPCR.**
(A) qPCR analysis confirming successful overexpression of MDM2 in U2OS cells following plasmid transfection for 24 h. (B) qPCR validation of miR-16-1-3p mimic overexpression in U2OS cells 24 h after transfection (50 nM). (C) qPCR analysis showing stable overexpression of miR-16-1-3p in U2OS cells generated by lentiviral infection and FACS sorting. Data are presented as mean ± SD; ***P < 0.001.
